# High-throughput transmission electron microscopy with automated serial sectioning

**DOI:** 10.1101/657346

**Authors:** Brett J. Graham, David Grant Colburn Hildebrand, Aaron T. Kuan, Jasper T. Maniates-Selvin, Logan A. Thomas, Brendan L. Shanny, Wei-Chung Allen Lee

## Abstract

Transmission electron microscopy (TEM) is an essential tool for studying cells and molecules. We present a tape-based, reel-to-reel pipeline that combines automated serial sectioning with automated high-throughput TEM imaging. This acquisition platform provides nanometer-resolution imaging at fast rates for a fraction of the cost of alternative approaches. We demonstrate the utility of this imaging platform for generating datasets of biological tissues with a focus on examining brain circuits.

## Introduction

For more than half a century, electron microscopy (EM) has allowed the highest resolution imaging of densely stained biological specimens^1–3^. Applied to the nervous system, EM allows unbiased labeling of fine neuronal processes (<200 nm), synaptic vesicles (∼40 nm), synaptic clefts (∼20 nm), and other intracellular components. By collecting, imaging, and aligning serial sections, volumetric datasets can be captured for comprehensive identification and classification of synapses and morphological categorization of cell types. However, even seemingly small tissue volumes (1 mm^3^) acquired at high resolution (e.g. 4 × 4 × 40 nm^3^ per voxel) produce massive datasets (>1500 teravoxels) that require automated methods for reliable acquisition in a reasonable amount of time. Recent developments in automated methods for scanning electron microscopy (SEM)—including serial block-face SEM (SBEM)^4^, focused ion beam milling SEM (FIB-SEM)^5,6^, and automated tape-collecting ultramicrotome SEM (ATUM-SEM)^7^—have enabled analysis of multiple neuronal circuits^8–16^. Transmission EM (TEM) allows for higher spatial resolution^17^, an order of magnitude greater signal-to-noise at the same electron dose^6,18^, and straightforward parallelization using fast off-the-shelf cameras^19–21^. However, development of automated methods for TEM has been limited (cf. ref. 18), in large part due to the lack of an automated sample collection, handling, and imaging pipeline compatible with transmitted electron detection. Here, we present an automated, tape-based data acquisition platform (Fig. 1) that combines automated sample collection with a novel TEM-compatible collection substrate and a reel-to-reel imaging stage that accelerates TEM imaging to >30 Mpixels per s at a relatively low cost (∼US$300,000 per microscope; see Table 1 and Supplementary Table 1).

**Table 1.** Serial-section EM microscope throughput and cost comparison. Based on published datasets: FIB-SEM resolution range: x,y,z: 5–8 nm^5,6^, SBEM: resolution ranges x,y: 9–16 nm, z: 20–30 nm^8,15,24^. For FIB-SEM+HotKnife^25^ imaging rates, K. Hayworth, personal communication; ATUM-MultiSEM resolution and imaging rates, R. Schalek, personal communication; and for TEMCA2-AutoLoader^18^ and C. Robinson, personal communication.

|  | **FIB-**<br>**SEM+HotKnife** | **SBEM** | **ATUM-**<br>**MultiSEM** | **TEMCA2-**<br>**AL** | **TEMCA-**<br>**GT** |
| --- | --- | --- | --- | --- | --- |
| x,y resolution (nm) | 8 | 9 | 4 | 4 | 4 |
| z resolution (nm) | 8 | 20 | 30 | 40 | 40 |
| *peak* imaging rate (Mpixels per s) | 8 | 0.5 | 250 | 50 | 80 |
| *effective* imaging rate (Mvoxels per s) | 2 | 0.2 | 130 | 6 | 30 |
| unattended operation | yes | yes | yes | yes | yes |
| microscope cost (each) | \$1M | \$1M | \$4-6M | \$1M | \$300k |
| **To image 1×0.5×0.5 mm<sup>3</sup> (cortical column)**<br>**on a single microscope** |  |  |  |  |  |
| Typical *effective* cutting rate (sections per day) | in effective<br>imaging rate | in effective<br>imaging rate | 2500 | 600 | 2500 |
| Number of sections | 125000 | 25000 | 16667 | 12500 | 12500 |
| Time to section (days) | n/a | n/a | 7 | 21 | 5 |
| Pixels per section | 3.91×10<sup>9</sup> | 6.17×10<sup>9</sup> | 3.13×10<sup>10</sup> | 3.13×10<sup>10</sup> | 3.13×10<sup>10</sup> |
| Effective time to image (days) | 2826 | 8564 | 46 | 768 | 151 |

**Figure 1.**
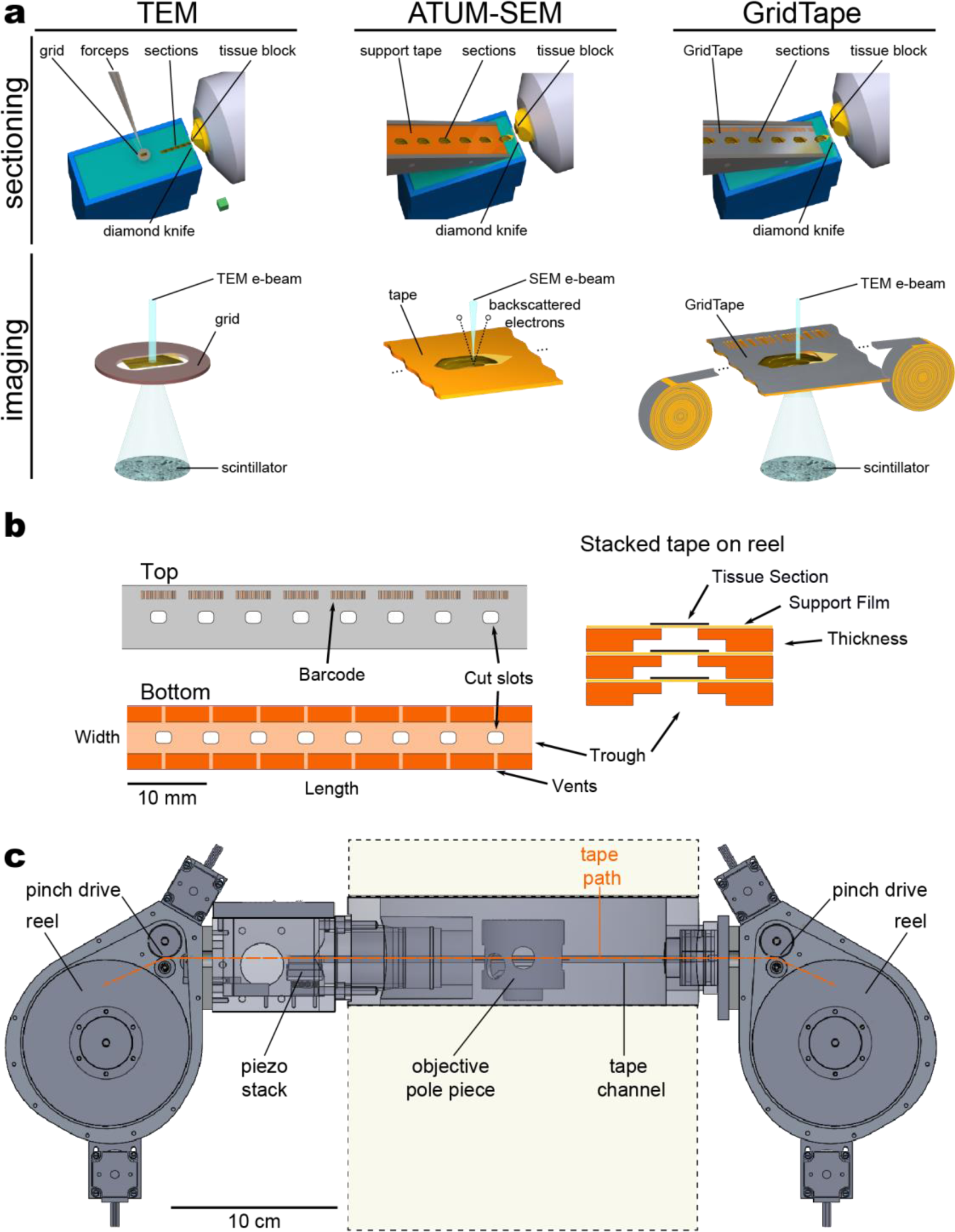
GridTape transmission electron microscopy (TEM) imaging platform. **a**, Schematic of (*left*) manual serial-section collection and TEM imaging; (*middle*) automated tape-collecting ultramicrotome (ATUM) collection and scanning electron microscopy (SEM) imaging, and (*right*) GridTape sectioning and TEM imaging. Ellipses denote continuation of tape. Bottom schematics not to scale. **b**, Schematic of the GridTape collection substrate (*left*) *en face* and (*right*) in cross-section. For clarity, tape thickness is exaggerated in the cross-section schematic and slot holes in different layers of tape are not necessarily aligned on top of one another. **c**, Schematic of the GridTape reel-to-reel stage at the level of the TEM objective pole-piece. Portions of the TEM column above and below in beige with dashed outlines. Scale box, 2 mm sides (**a**(*top*)); scale bars, 10 mm (**b**(*left*)) and 10 cm (**c**).

## Results

We developed a TEM-compatible tape substrate, called ‘GridTape’ (Fig. 1a–b), that combines advantages of automated section collection from ATUM-SEM^7^ with high-throughput TEM imaging^19–21^ (Fig. 1a). To produce GridTape, regularly spaced holes (resembling slots in conventional TEM grids; Fig. 1b) were laser milled through aluminum-coated polyimide (Kapton®) tape. The cut tape was then coated with a 50 nm-thick electron-lucent film (LUXFilm®) that spanned the holes and provided support for subsequent section collection. We collected sections onto GridTape with a conventional ultramicrotome and modified ATUM (Supplementary Fig. 1 and Methods). Resin-embedded samples were trimmed with tapered leading and trailing edges (Figs. 1a, 2c,f,i) and the ATUM was positioned close to the knife edge. This permitted collection of sections directly onto the moving tape, which facilitated consistent section placement. By monitoring the ultramicrotome cutting arm and adjusting the speed of the tape, the movement of GridTape slots was locked in-phase with cutting. This closed-loop sectioning approach enabled automated collection of >4000 sections per day with reliable positioning of sections over the slots.

To image sections collected on GridTape, we engineered an automated reel-to-reel TEM sample stage (Fig. 1c). The stage was mounted to standard X-ray detector ports with feed and pickup reel housings on opposite sides of the TEM column. Reel and pinch drive motors advanced the tape until the target section was positioned under the electron beam. The tape was then rapidly translated using piezoelectric nano-positioners while a camera array captured wide-field images to montage the region of interest.

We next assessed the capability of a TEM camera array (TEMCA)^19–21^ combined with GridTape (TEMCA-GT) to generate large serial-section EM datasets of tissue volumes. As an initial test, we produced a 5.6 million µm^3^ dataset of the mouse dorsal lateral geniculate nucleus (dLGN) consisting of 250 serial sections (Fig. 2a–c and Supplementary Table 2). The sections were collected on top of the coated holes of the GridTape with high precision (Fig. 2c,j; 98% < 0.26 mm positional deviation) and no sections were lost. We then acquired a region of interest from the series on one TEMCA-GT at an imaging rate of 38.01 ± 0.19 (mean ± SE) Mpixels per s. As with all other datasets described here, each section was acquired with 4.3 nm per pixel lateral resolution. Individual sections spanned the dorsoventral and mediolateral extent of the dLGN, including the core and shell regions, and synaptic connections were resolvable (Fig. 2a–b). Retinal and cortical inputs to the dLGN were readily distinguishable based on ultrastructural features including the presence of large, pale mitochondria.

**Figure 2.**
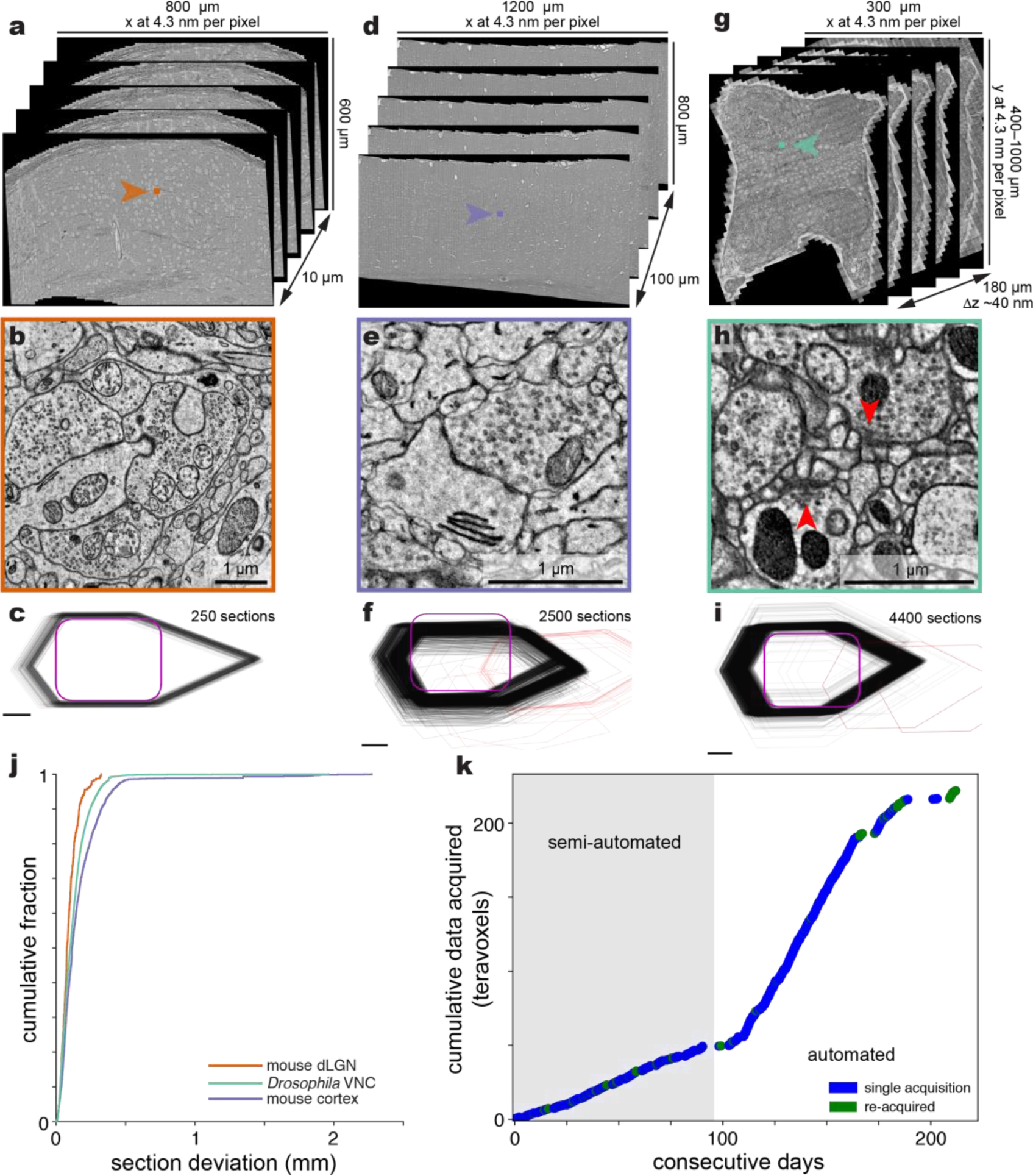
Volumetric EM datasets acquired using GridTape. Data from (**a**–**c**) mouse dorsal lateral geniculate nucleus (dLGN), (**d**–**f**) mouse cortex, and (**g**–**i**) *Drosophila* ventral nerve cord (VNC) series. **a**, Exploded, zoomed-out views of dLGN EM volume. **b**, Zoomed-in view of synaptic contacts from the boxed region in (**a**). **c**, Positions of all sections in the dLGN series (elongated hexagonal outlines) relative to their slot (purple outline). **d**, Exploded, zoomed-out views of cortex EM volume. **e**, Zoomed-in view of a synapse from the boxed region in (**d**). **f**, As in (**c**), for cortex sections. Red outlines indicate sections where the region of interest was fully off the slot and could not be acquired with TEM. **g**, Exploded, zoomed-out views of VNC EM volume. **h**, Zoomed-in view of synaptic contacts from the region in (**g**). Red arrowheads indicate pre-synaptic specializations known as T-bars. **i**, As in (**c**), for VNC sections. **j**, Cumulative distribution of section placement deviation (Euclidean distance from the average section position) for each dataset. **k**, TEMCA-GT imaging rate of the mouse cortex volume expressed as cumulative data acquired over consecutive days. Colors represent whether sections were re-acquired to improve focus or region of interest placement. The dLGN and VNC datasets were acquired almost entirely in automated mode. Scale bars, 1 µm (**b, e, h**) and 500 µm (**c, f, i**).

We then scaled up to generate datasets spanning thousands of serial sections from the mouse cortex (Fig. 2d–f and Supplementary Table 2) and *Drosophila* ventral nerve cord (VNC) (Fig. 2g–i and Supplementary Table 2). We produced a 225 trillion voxel dataset from the mouse cortex spanning 166 million µm^3^ (Fig. 2d–f). Most sections (98% of 2500) were positioned within 0.49 mm from the average section position (Fig. 2f,j) with 62 of 2500 sections lost. With 2 mm-long slots, this allowed for reliable imaging of regions at least 1 mm long. After transitioning to automated TEMCA-GT imaging both within and across sections, we imaged the last 2167 sections for this cortical dataset at a rate of 39.59 ± 0.08 (mean ± SE) Mpixels per s (Fig. 2k).

Finally, we generated an 86.7 trillion voxel dataset spanning 64 million µm^3^ of an adult female *Drosophila* VNC cut horizontally (Fig. 2g–i and Supplementary Table 2). Most sections (98% of 4400) were positioned within 0.37 mm from the average section position (Fig. 2i–j) with 5 of 4400 sections lost. The VNC dataset was imaged at a rate of 42.73 ± 0.05 (mean ± SE) Mpixels per s. Together, our results demonstrate the general ability of collecting thousands of serial sections onto a tape-based substrate and imaging them at nanometer resolution reliably and rapidly with TEM.

## Discussion

Development of an automated, reel-to-reel pipeline for TEM imaging further underscores TEM’s vital role in scalable, high-resolution structural imaging. With the potential of efforts such as EM connectomics, multiple techniques for data collection are key to safeguard against unforeseen future challenges. Each large-scale EM approach^4,6,7,19^ has its own advantages. We propose GridTape as a complementary approach. Combined with TEMCA imaging^19–21^, it provides a workflow that has advantages of high signal-to-noise and is inherently parallel. This results in imaging speed and scalability, with lower capital costs. For the current price of one commercial multi-beam SEM system, ten TEMCA-GTs can be built and samples collected on tape can be distributed across machines for simultaneous imaging. Moreover, acquisition rates will further increase as cameras and imaging sensors continue to improve. The GridTape approach also has limitations. Imaging area is currently constrained by slot size. Although features milled in the tape are customizable, imaging regions will ultimately be limited by microscope geometry and material properties of the thin-film coating. Additionally, the fixed costs for the microscope hardware are accompanied by consumable costs associated with the tape and its thin-film coating (currently ∼USD$4 per slot). However, we expect these costs to decrease with GridTape adoption due to economies of scale.

Because samples are not destroyed during imaging, large-scale, serial-section TEM can be enhanced with post-section labeling and combined with optical imaging. More broadly, as ever larger sample regions can be simultaneously imaged at high resolution, throughput becomes ultimately limited by the speed and precision of sample movement. Automated, continuous, reel-to-reel sample handling on transmissive substrates may offer a general approach for high-throughput data collection across multiple imaging modalities.

The GridTape approach provides accessible, high-throughput imaging that will be useful for studies of development, examining the variability between individuals, sexually dimorphic features, whole brains or organisms, and to study results of experimental perturbations or mutant phenotypes. Considered more generally, it serves as an advance in the rich history of EM, particularly benefitting applications that leverage the advantages of transmitted electron detection.

## Acknowledgements

We thank C. Bolger and the HMS EM Core Facility for technical support; X. Chen and R. Zheng for programming and software support; A. Bleckert, D. Brittain, R. Torres, N. da Costa, and R.C. Reid at the Allen Institute for Brain Science for their encouragement, feedback, and support; K. Hayworth for his vision, inspiration, and advice; J. Lichtman, R. Schalek, and K. Hayworth for an ATUM device; O. Mazor and P. Gorelik for engineering support; R. Fetter for histology advice; C. Guo, W. Regehr, L. Cheadle, M. Greenberg, L. Driscoll, and C. Harvey for generously donating samples; N. Perrimon for generously sharing *Drosophila* lines before their publication; S. Rayshubskiy and R. Wilson for sharing flies; H. Somhegyi for assistance with *Drosophila* dissections; T. Ayers, R. Smith, and Luxel Corporation for coating tape; and L. Cheadle, D. Ginty, C. Harvey, W. Regehr, E. Soucy, and the Lee laboratory for comments on the manuscript. This work was supported by NIH grants (R21NS085320, RF1MH114047) to W-C.A.L., the Bertarelli Program in Translational Neuroscience and Neuroengineering, Edward R. and Anne G. Lefler Center, Stanley and Theodora Feldberg Fund, and the Intelligence Advanced Research Projects Activity (IARPA) of the Department of Interior/Interior Business Center (DoI/IBC) through contract number D16PC00004; and by NIH grants (T32MH20017, T32HL007901) for D.G.C.H. Portions of this research were conducted on the Orchestra High Performance Compute Cluster at Harvard Medical School partially provided through NIH NCRR grant 1S10RR028832-01. The views and conclusions contained herein are those of the authors and should not be interpreted as representing the official policies or endorsements, either expressed or implied, of the funding sources including NIH, IARPA, DoI/IBC, or the U.S. Government.

## Author contributions

D.G.C.H. and W-C.A.L. conceptualized the project. B.J.G. and D.G.C.H. designed, built, and developed software for tape milling. B.J.G, D.G.C.H and B.L.S. designed, built, and developed software for the ATUM and ultramicrotome modifications. B.J.G. designed, built, and developed software for tape handling, computerized microscope control, and the reel-to-reel GridTape imaging stage. B.L.S. developed software for tape staining and the reel-to-reel GridTape imaging stage. B.J.G., D.G.C.H., B.L.S., J.M-S., L.A.T., and A.T.K. developed control and analysis software. J.M-S. and W-C.A.L. prepared samples. D.G.C.H., J.M-S., and A.T.K. sectioned samples. B.J.G., D.G.C.H., B.L.S., J.M-S., L.A.T., A.T.K., and W-C.A.L. collected data. B.J.G., J.M-S., L.A.T., and A.T.K. stitched and aligned datasets. B.J.G., D.G.C.H., and W-C.A.L. wrote the paper.

## Methods

### Animals and tissue preparation

All procedures involving animals were conducted in accordance with the ethical guidelines of the NIH and approved by the IACUC at Harvard Medical School. The Standing Committee on the Use of Animals in Research and Training of Harvard University approved all animal experiments. In most cases, mammalian tissue was collected from animals that were previously participants in other experiments.

#### Mouse cortex

We prepared one mouse cortex specimen for EM as described previously^19,20^. A 5-month-old male C57BL/6J mouse was perfused *trans*-cardially (2% formaldehyde and 2.5% glutaraldehyde in 0.1 M cacodylate buffer with 0.04% CaCl_2_) following *in vivo* two-photon imaging, and the brain was processed for serial-section TEM. Briefly, 300 µm-thick coronal vibratome sections were cut, post-fixed, and *en bloc* stained with 1% osmium tetroxide and 1.5% potassium ferrocyanide followed by 1% uranyl acetate, followed by lead aspartate, dehydrated with a graded ethanol series, and embedded in epoxy resin (TAAB 812 Epon, Canemco). Following section collection onto GridTape (see ‘Serial sectioning’), sections were stained with 3% lead citrate (Ultrostain II, Leica) for 5 minutes (see ‘Post-section staining’).

#### Mouse dLGN

One mouse dLGN specimen was prepared with enhanced *en bloc* staining based on previously described protocols^22,23^. The specimen was from a P27 male C57BL/6J mouse. Following section collection onto GridTape, sections were each stained with 3% lead citrate (Ultrostain II, Leica) for 5 minutes (see ‘Post-section staining’).

#### Drosophila

We fixed one adult female *Drosophila melanogaster* (aged 1 day post-eclosion, genotype VT25718-Gal4/+; UAS–3×MYC–sbAPEX2–dlg/+) specimen as described previously^21^; stained with enhanced *en bloc* staining^22,23^; and embedded the central nervous system in epoxy resin, positioned in a cutout of EM-prepared mouse cortex^16^. Sections were not post-section stained.

### Substrate production

GridTape was produced from 125 µm aluminum-coated Kapton® film (Dunmore) slit into 8 mm-wide reels of over 35 m in length (Metlon Corporation). This stock tape was modified using a custom laser-milling system consisting of a reel-to-reel tape positioning machine and commercial 1 W ultraviolet laser marking system (Samurai, DPSS Lasers). Control software triggered laser milling of a 30 mm length of tape, used custom computer vision to check the result of the cutting, advanced the tape 30 mm and finally adjusted the position of the tape to align the next 30 mm of tape to cut. This system enabled the autonomous production of >30 m lengths of cut tape containing over 5000 slots. Following laser milling, the cut tape was cleaned by wiping it with isopropyl alcohol-soaked lint free wipes. Finally, the cut tape was sent to Luxel Corporation for application of a 50 nm TEM support film (LUXFilm®).

### Sample block trimming

In preparation for sectioning, embedded tissue blocks were trimmed (Trim 90, Diatome) into an oblong hexagonal shape (Fig. 2c,f,i) with 3.5–4 mm height, 1–2 mm width, a greater than 90° degree bottom angle and less than 90° top angle.

### Serial sectioning

An ultramicrotome (UC7, Leica) and diamond knife (4 mm, 35° Ultra or Ultra-Maxi, Diatome) was used to cut ultra-thin serial sections (∼40 nm) from prepared samples. These sections were collected using a modified automated tape-collecting ultramicrotome (ATUM)^7^. All tape guides and rollers on the ATUM were modified by adding a 4 mm-wide trough to prevent contact with the TEM support film spanning GridTape slots. Additionally, an optical interrupter (GP1A57HRJ00F, Sharp) was affixed to the ATUM to detect the passage of GridTape slots and a hall-effect sensor (A1301EUA-T, Allegro MicroSystems) and magnet were attached to the microtome swing arm to detect the cutting of sections. Custom software monitored the period and relative phase-offset of these two sensors during section collection. By setting the microtome to a fixed cutting speed and varying the ATUM tape speed, effective phase-locking at a fixed offset was achieved. This system (Supplementary Fig. 1) provided accurate placement of cut sections atop GridTape slots (Fig. 2c,f,i,j).

### Measuring section placement consistency

Section placement was measured by first capturing light-level images (Flea3 FL3-U3-13E4C-C, PointGrey) of each target slot with collimated low-angle illumination (MWWHL4, Thorlabs). Slot and section perimeters were located manually by adjusting the position and rotation of a template outline for each feature to each section using a custom MATLAB (MathWorks) script. Although this process could be accurately performed in tens of seconds per section, an automated routine was also used to speed up annotation of large datasets. For the automated approach, images were processed using the ImageJ plugin “Template Matching and Slice Alignment” (https://sites.google.com/site/qingzongtseng/template-matching-ij-plugin) to find the location of each slot and corresponding section. The plugin correctly identified the location of the slot in >99% of images by finding the image region that most resembled a template image of a slot. Any errors were corrected manually in ImageJ. Next, to measure the offset from the slot to the tissue section, a prominent feature of the tissue section was chosen as a template. The plugin correctly identified the location of that tissue feature in ∼98% of images. This success rate was achieved by choosing a small, high-contrast feature of the tissue that tended to be placed close to the middle of the slot. For series where the tissue features significantly changed shape over the course of the pickup, template matching was performed on smaller batches of 500–1000 sections, with a separate feature chosen for template matching in each batch. Finally, the remaining ∼2% of sections that were not correctly located with the template matching plugin were located manually in ImageJ. The sections needing this manual correction mainly fell into two categories: sections that were cut very thin, causing the tissue features to have reduced contrast, or sections with the template feature placed off-slot.

### Post-section staining

To enhance sample contrast, post-section staining was performed on most samples by application of Reynolds lead citrate (Ultrostain II, Leica). A benchtop reel-to-reel system was constructed that exposed ∼50 slots of GridTape spanned between two controllable pinch drives. A droplet of stain was manually applied to a subset of exposed GridTape slots. After incubation (typically 5 min) an array of nozzles were used to wash double-distilled water over the surface of the GridTape for 1 min. During this wash period, a matching array of suction nozzles prevented overflow.

### TEM imaging

To perform TEM GridTape imaging, a custom in-vacuum, reel-to-reel stage was constructed (Fig. 1c). This stage allows a full 45 m-long roll of GridTape to be loaded into the microscope for imaging. After loading and pump-down, a set of pinch drives (one on each side of the TEM column) allows linear movement of GridTape to exchange and position sections under the beam in preparation for imaging. After positioning, both pinch drives dispense a small amount of GridTape towards the center of the column, introducing slack on both sides of the sample held under the beam. This allows an XY stack of piezo nanopositioners (SLC-1720, SmarAct) to make the many small movements necessary to montage large areas. At 4.3 nm lateral resolution, the TEMCA field of view for a single location was just over 16µm square. By capturing many images at slightly overlapping regions (typically 20–30%) for a single section, regions of interest in excess of 1 mm^2^ were imaged. After acquisition, camera images were virtually stitched and aligned with AlignTK (http://mmbios.org/installation) into three-dimensional (3-D) volumes^19–21^.

## Data availability

GridTape instrumentation designs will be made publicly available upon peer-reviewed publication of this pre-print. The image datasets generated and analyzed for this study will be made available through separate publications. In the interim, to request access to data, please contact the corresponding authors.

## Code availability

Custom software tools generated for data handling, visualization, and analysis are available from the corresponding authors upon reasonable request.

**Supplementary Table 1.**
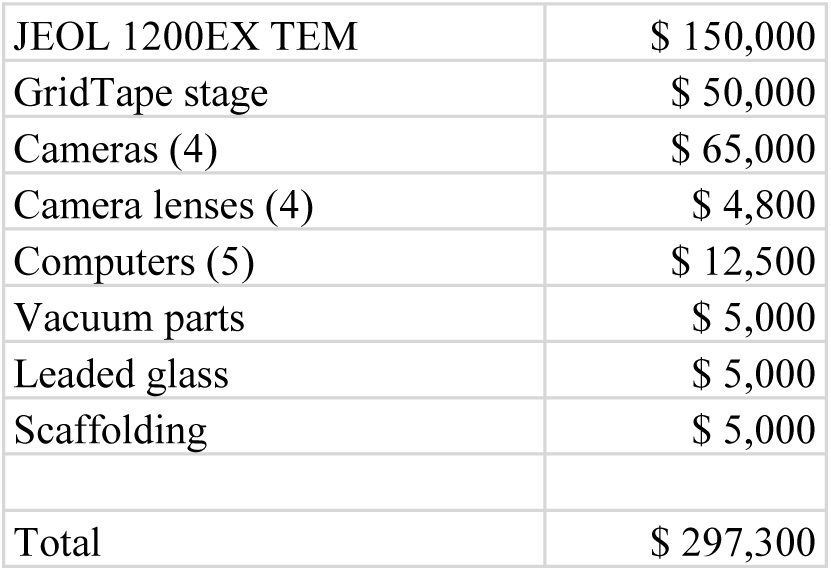
Approximate TEMCA-GT cost.

**Supplementary Table 2.** Automated TEMCA-GT dataset metrics: ‘active imaging rate’ is the rate during image acquisition whereas ‘imaging rate with overhead’ also includes time moving between sections and imaging setup (i.e. electron beam adjustment, sample pre-bake, and focus).

| dataset | size<br>(Tvoxels) | size<br>($\mu\text{m}^3$) | active rate<br>(Mpix/s) ± S.E.M. | rate with overhead<br>(Mpix/s) ± S.E.M. | sections manual<br>intervention (%) | sections<br>reimaged (%) |
| --- | --- | --- | --- | --- | --- | --- |
| mouse dLGN | 7.5 | $5.55 \times 10^6$ | $38.01 \pm 0.19$ | $37.70 \pm 0.90$ | 6.4 | 1.2 |
| mouse cortex<br>(*automated*<br>*portion only*) | 130 | $9.60 \times 10^7$ | $39.59 \pm 0.08$ | $37.04 \pm 0.23$ | 9.9 | 6.2 |
| *Drosophila*<br>VNC | 86.7 | $6.41 \times 10^7$ | $42.73 \pm 0.05$ | $26.10 \pm 0.10$ | 2.2 | 5.5 |

**Supplementary Figure 1.**
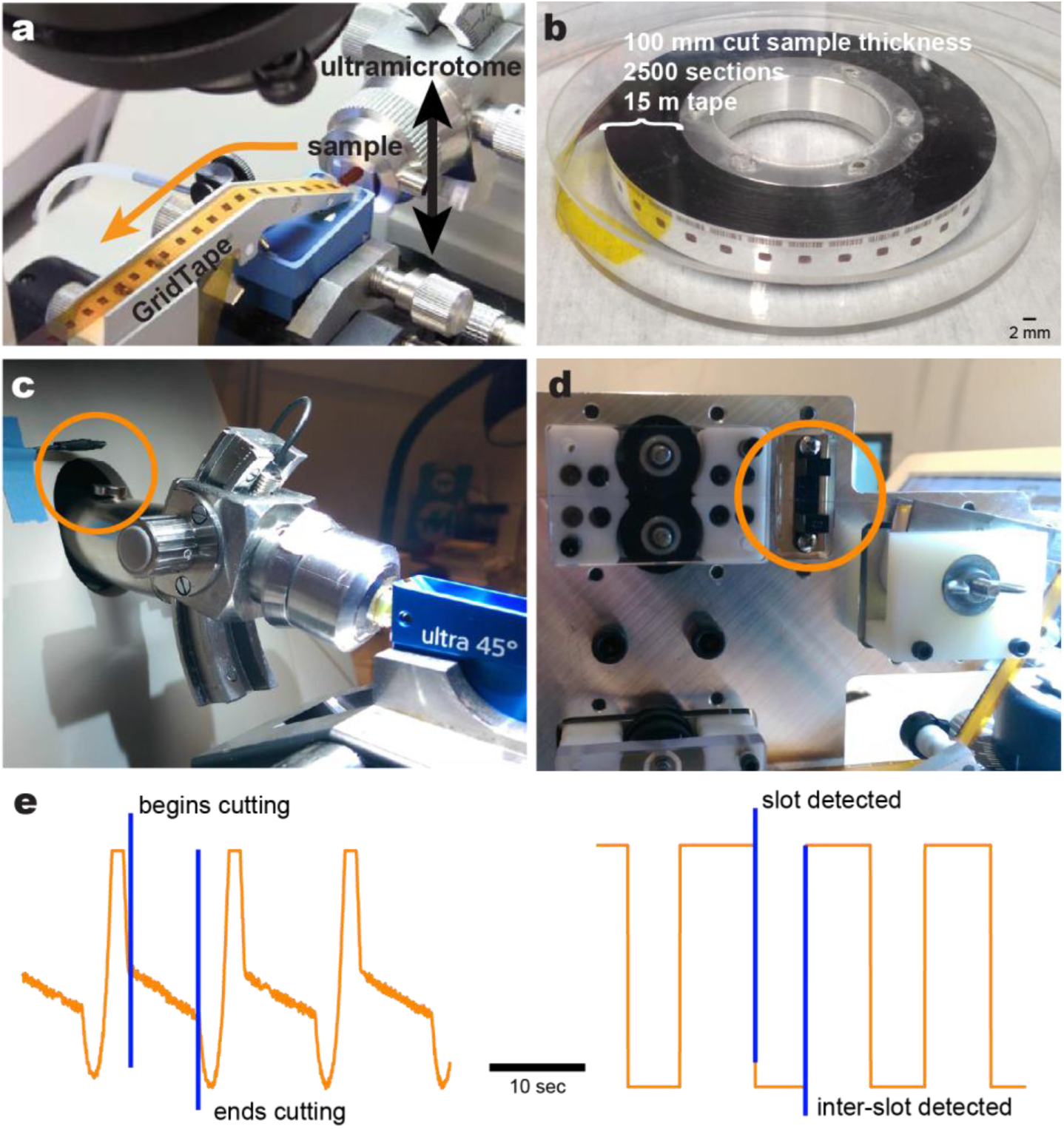
GridTape section collection. Photographs of **a**, the ATUM collection system with sample sections (dark rectangles) collected on top of non-aluminum coated GridTape (yellow-orange), the diamond knife boat (blue), and the sample mounted at the end of the ultramircotome cutting arm. **b**, a 15 m long reel of GridTape holds 2500 serial sections or 100 µm of sample thickness cut at 40 nm thin sections. **c**, the magnet and hall effect sensor (circle) attached to the microtome cutting arm to measure cutting frequency. **d**, the digital opto-interrupter (circle) used to detect the slot frequency in the tape. **e**, The (*left*) analog signal from the hall effect sensor and (*right*) digital signal from the opto-interruper are used to perform closed-loop, phase-locked sectioning. Scale bar, 2 mm (**b**).

